# Systematic part transfer by extending a modular toolkit to diverse bacteria

**DOI:** 10.1101/2023.02.07.527528

**Authors:** Kevin Keating, Eric M. Young

## Abstract

It is impractical to develop a new parts collection for every potential host organism. It is well-established that gene expression parts, like genes, are qualitatively transferable, but there is little quantitative information defining transferability. Here, we systematically quantified the behavior of a parts set across multiple hosts. To do this, we developed a broad host range (BHR) plasmid system compatible with the large, modular CIDAR parts collection for *E. coli*. This enabled testing of a library of DNA constructs across the Pseudomonadota – *Escherichia coli, Pseudomonas putida, Cupriavidus necator*, and *Komagataeibacter nataicola*. Part performance was evaluated with a standardized characterization procedure that quantified expression in terms of molecules of equivalent fluorescein (MEFL), an objective unit of measure. The results showed that the CIDAR parts enable graded gene expression across all organisms – meaning that the same parts can be used to program *E. coli, P. putida, C. necator*, and *K. nataicola*. Most parts had a similar expression trend across hosts, although each organism had a different average gene expression level. The variability is enough that to achieve the same MEFL in a different organism, a lookup table is required to translate a design from one host to another. To identify truly divergent parts, we applied linear regression to a combinatorial set of promoters and ribosome binding sites, finding that the promoter J23100 behaves very differently in *K. nataicola* than in the other hosts. Thus, it is now possible to evaluate any CIDAR compatible part in three other hosts of interest, and the diversity of these hosts implies that the collection will also be compatible with many other Proteobacteria (Pseudomonadota). Furthermore, this work defines an approach to generalize modular synthetic biology parts sets beyond a single host, making it possible to create a small number of parts sets that can span the tree of life. This will accelerate current efforts to engineer diverse species for environmental, biotechnological, and health applications.

## INTRODUCTION

Bacteria with powerful metabolic capabilities that are well-adapted to challenging environments promise solutions for a variety of current manufacturing and environmental problems^1^. Tools for genetic manipulation of these organisms are required to study their phenotypes and design them to achieve set objectives. To this end, synthetic biology and metabolic engineering are currently engaged in efforts termed “onboarding” that seek to optimize and accelerate successful genetic transformation and genome engineering across the tree of life. Though new toolkits are continually published for organisms across the bacterial domain^2–10^, these toolkits often focus on single species or a narrow range of related bacteria with attractive catabolic, anabolic, and tolerance phenotypes. However, this approach will struggle to keep pace as the breadth and scale of engineerable microbes increases. Ideally, designers should be able to reprogram organisms as quickly as they are described. Therefore, it is not practical to develop new tools for each organism. Instead, methods for transferring genetic parts between hosts are needed.

Part transfer is common in nature. Genomics has confirmed horizontal gene transfer (HGT) between organisms, which can confer a competitive advantage to the new host^11^. Mobile bacterial genetic elements, including plasmids and transposons^12^, also commonly have broad host ranges (BHR). To replicate within the host or to provide an evolutionary advantage, these elements carry genes for cisregulatory proteins, antibiotic resistance, biosynthetic gene clusters, or a variety of other functions. The genes are expressed, implying that either leaky transcription or the original gene expression parts are able to transcribe the foreign genes. These elements have been repurposed into vectors since the beginning of genetic engineering, and continue to form the foundation for new broad host range expression strategies^13,14^. It is clear from these examples that functional part transfer should be possible, but how broad, quantifiable, and predictable genetic part transfer can be has not yet been established.

Paired promoters and ribosome binding sites (RBS) are elemental genetic parts in gene expression control. Synthetic biology has widely adopted the Anderson promoter library, a series of synthetic constitutive promoters designed as variants of a consensus *E*.*coli* promoter, and synthetic RBSs from the Weiss group and the BIOFAB Foundry to achieve graded constitutive gene expression in *E. coli*^5,15^. Recently, these parts were incorporated into a modular cloning standard, CIDAR, that permits combinatorial assembly. This standard has been recently expanded to include over 100 unique parts, making it the largest of the *E. coli* part kits available on Addgene. Many of these parts have been used in a range of bacteria outside of *E. coli*^4,16^, yet their function has not been systematically characterized between diverse species. Often, the plasmid backbone, part context, or flow cytometry method varies between these experiments. Therefore, it is difficult to compare expression across these instances because the same DNA and quantification methods have not been used.

The phylum *Pseudomonadota* contains several species of interest that could be new hosts for synthetic biology, yet the available parts are limited. These species span the major classes within the phylum. *Pseudomonas putida*, like *E. coli*, is a gammaproteobacterium commonly found in soil that can degrade organic solvents, thrive in harsh chemical environments, and is genetically tractable with a few emerging parts sets^17^. The betaproteobacterium *Cupriavidus necator* is well-known for its flexible metabolism and high accumulation of polyhydroxybutyrates, yet there are few genetic tools available^18^. The alphaproteobacterium *Komagataeibacter nataicola*, along with related cellulose-producing acetic acid bacteria, is used in the fermentation of kombucha tea as well as the industrial production of bacterial nanocellulose^19^. Some engineering tools have been established in all of these organisms, but not at the scale and depth of the CIDAR parts for *E. coli*.

Here, we re-design the CIDAR backbones to include a broad host range replicon, BBR1, to enable CIDAR parts expression across *Pseudomonadota*. These backbones, termed openCIDAR, are compatible with all the parts currently on Addgene. We then characterize standard genetic parts in the lab strain *Pseudomonas putida* KT2440, the lab strain *Cupriavidus necator* H16, and a newly isolated strain of *Komagataeibacter nataicola* called DS12. This spans the major groups of the *Pseudomonadota* as well as strains in various stages of domestication. We then assess the function of CIDAR promoters and RBSs in the same genetic context in each of these hosts, using molecules of equivalent flourescein (MEFL), an emerging standard unit for flow cytometry^20,21^, to quantitatively compare performance. We find that the same DNA functions relatively the same across the hosts, showing that just as genes can be relatively easily exchanged across organisms, so can transcriptional and translational machinery. We also show that linear models can identify parts that behave less predictably, specifically J23100 in *K. nataicola*, enabling rapid identification of the appropriate parts to use in engineering a new host. Thus, a large, well-characterized parts set with graded gene expression is now accessible for any organism compatible with the pBBR1 plasmid, which is most *Pseudomonadota*.

## RESULTS

### Genetic design strategy

We wanted to design our system to be compatible with accessible commonly used parts. Therefore, we sought an assembly standard that had many well-characterized physical constructs readily available to researchers. We found that the Modular Cloning toolkit on Addgene that conforms to the CIDAR standard from the Densmore and Freemont labs, when combined with the extension kit available from the Murray lab, is comprehensive, characterized, accessible, and convenient to clone. All together, the CIDAR standard currently contains 15 constitutive promotes, 15 inducible promoters, 5 monocistronic RBSs, 14 bicistronic RBSs, 18 fluorescent protein coding sequences, 59 other coding sequences, and 11 terminators. Other possibilities like the iGEM collection, the DOE JGI Inventory of Composable Elements, or SynBioHub, met most, but not all, of our initial requirements. Therefore, everything we built in this effort is compatible with the CIDAR standard.

We next designed an expression vector based on a broad host range (BHR) plasmid. BHR plasmids are independent of host genomics and simple in terms of cloning. We considered this superior to integrative vectors, which are stable, yet they tend integrate randomly or require pre-insertion of a landing pad, creating context effects and increasing host engineering effort. There are several BHR plasmids that function in *Pseudomonadota* (Table 1). Of these, the pBBR1 plasmid is small, has a broad range, relatively high stability, and is maintained at a medium copy number (Figure 1A). In pBBR1, the *Rep* protein is required for replication, and in all known cases expression is controlled by DNA elements (*i*.*e*. promoter and RBS) native to the pBBR1 plasmid (Figure 1B). Thus, a backbone with a replicon based on BBR1 and compatible with the CIDAR Modular Cloning standard could extend well known synthetic biology parts across a wide range of bacterial hosts (Figure 1C). To test this, we selected four hosts (highlighted, Figure 1A) representing the major classes of *Pseudomonadota*.

**Table 1:**
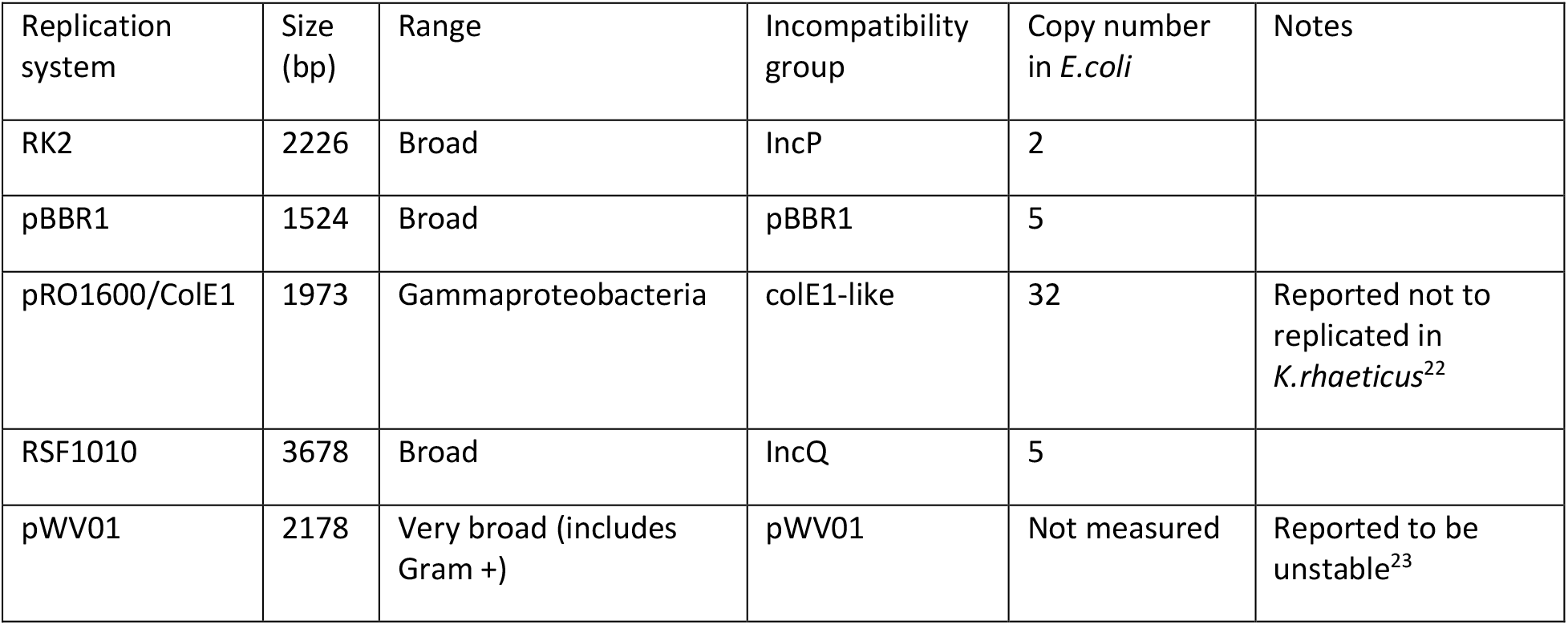
Broad host range plasmids vary in size, range, and copy number.

| Replication system | Size (bp) | Range | Incompatibility group | Copy number in *E. coli* | Notes |
| --- | --- | --- | --- | --- | --- |
| RK2 | 2226 | Broad | IncP | 2 |  |
| pBBR1 | 1524 | Broad | pBBR1 | 5 |  |
| pRO1600/ColE1 | 1973 | Gammaproteobacteria | colE1-like | 32 | Reported not to replicated in *K.rhaeticus*<sup>22</sup> |
| RSF1010 | 3678 | Broad | IncQ | 5 |  |
| pWV01 | 2178 | Very broad (includes Gram +) | pWV01 | Not measured | Reported to be unstable<sup>23</sup> |

**Figure 1:**
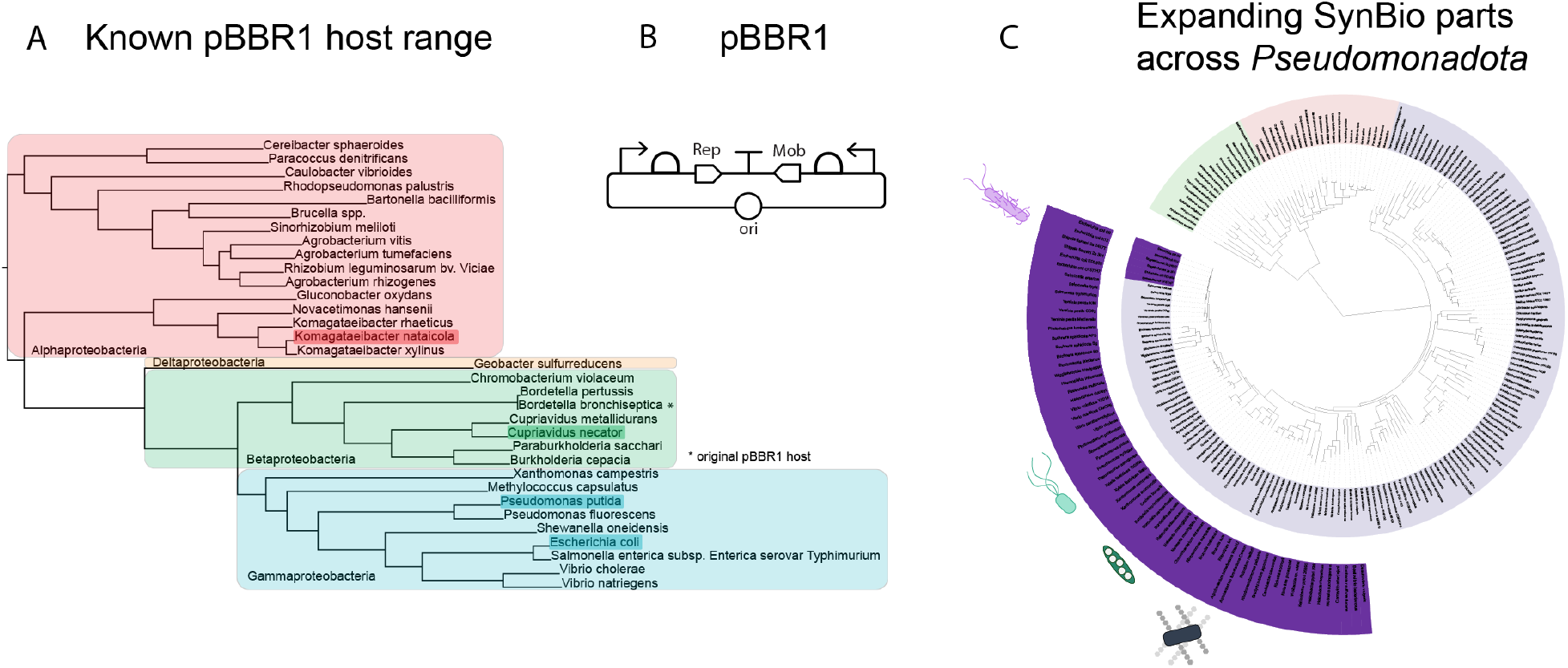
Enabling parts expression across hosts. **A**. Known pBBR1 compatible bacteria, presented in a reconstructed phylogenetic tree. All major clades of *Pseudomonadota* are represented in this group. Hosts used in this study are highlighted. **B**. Diagram of the native architecture of pBBR1, as isolated from *Bordatella bronchiseptica*. Modern derivatives do not include the *Mob* transcription unit, but are otherwise similar. **C**. The tree of life. The current host range of *E. coli* synthetic biology parts are highlighted within the inner arc. However, given the known range of pBBR1, the possible range is actually the outer highlighted arc.

### Creating openCIDAR – a broad host range parts collection

First, we built destination vectors that combine the CIDAR standard with the pBBR1 origin of replication. In order to be compatible, a synonymous mutation was introduced to the *rep* protein to remove a BbsI site. The resulting vectors are listed in **Error! Reference source not found**.. For the Level 1 destination vectors, we retained the CIDAR kanamycin marker for selection because the target species in this study are sensitive to kanamycin (**Error! Reference source not found**.)^16,24^. However, we replaced the Level 2 ampicillin marker with spectinomycin and chloramphenicol because *K. nataicola* was found to be only moderately sensitive to ampicillin (**Error! Reference source not found**.).

Next, we characterized the copy number of the new vectors in each host organism, to confirm the mutated pBBR1 was still functional. The Level 1 (single transcription unit) destination vector pBBR1_DVK_AE was transformed into all host strains, and plasmid copy number during exponential growth phase was determined by ddPCR. Copy number was defined as the ratio of absolute concentration of BBR1 *rep* protein amplicons to a single-copy target in each host genome (**Error! Reference source not found**.). The copy numbers for *E. coli* and *P. putida* were similar, with approximately 10-11 copies per cell for each species (Figure 2C and D). The average copy number was lower for *C. necator* and *K. nataicola*, at about 7 (Figure 2E) and 3 (Figure 2F) copies per cell, respectively.

**Figure 2:**
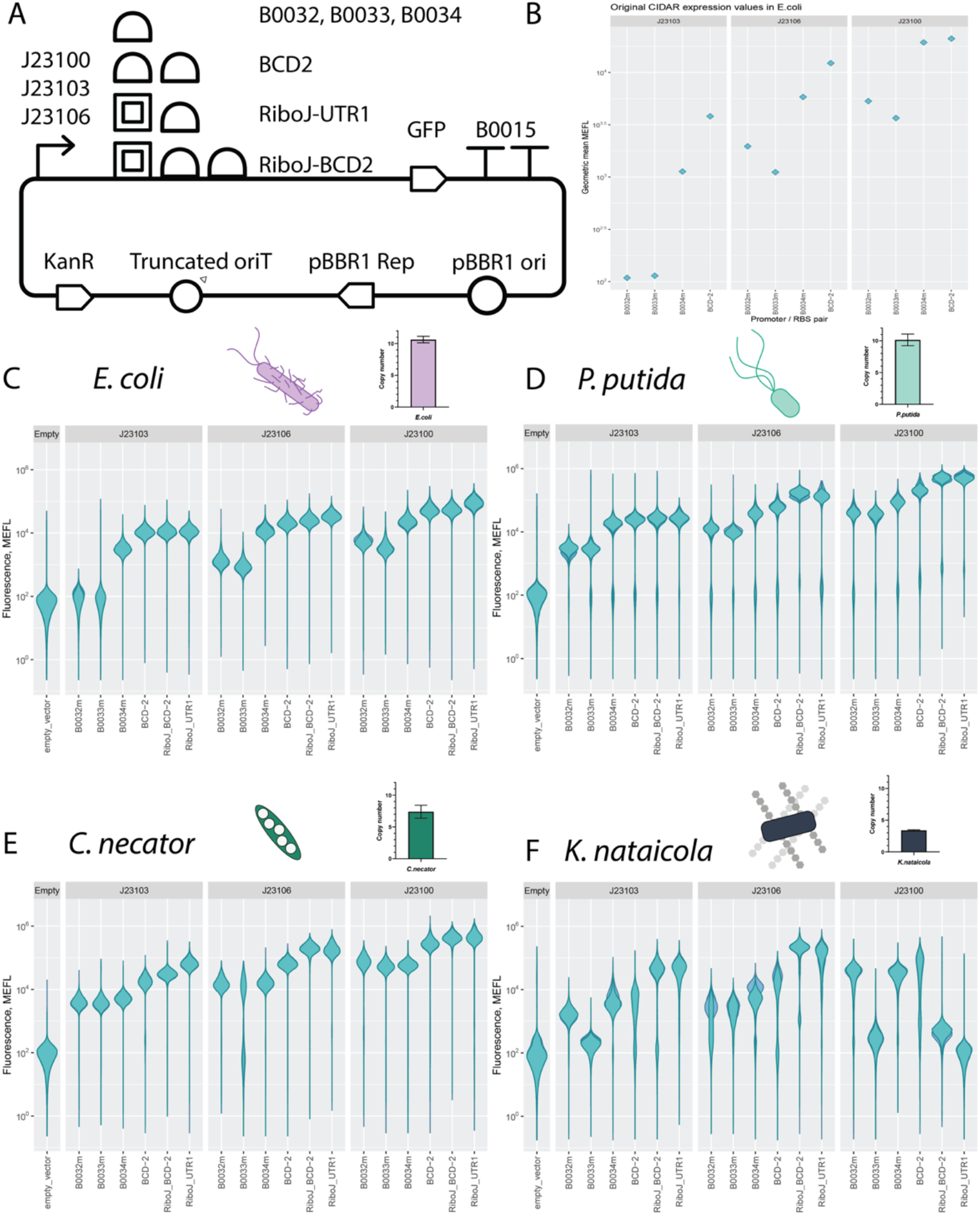
openCIDAR design and function. **A**. Design of the new destination vectors containing KanR, the pBBR1 origin and *rep* gene, and a CIDAR cloning site where transcription units may be assembled. Although this is over 100 parts, only the parts in this study are shown. **B**. Molecules of Equivalent Fluorescein (MEFL) expression values for the combinations of parts included in the original CIDAR paper. These were interpolated from Supplementary Figure 1. Expression from the original vector series in *E*.*coli* matches closely to the expression reported in this study. **C**. Violin plots of the expression values in MEFL for *E. coli* populations expressing each construct, along with the average copy number assessed by ddPCR. The violin plots for three replicates are overlaid, showing little variation between replicates. **D**. The same plots in as in C for the same constructs expressed in *P. putida*. **E**. The same plots in as in C for the same constructs expressed in *C. necator*. **F**. The same plots in as in C for the same constructs expressed in *K. nataicola*.

### Analysis of a combinatorial expression library demonstrates wide portability of commonly-used parts

To test gene expression across the hosts with the openCIDAR vectors, we selected commonly used parts that have been well characterized in *E. coli* (Figure 2A). For promoters, synthetic Anderson series promoters were chosen. In *E. coli* the expected qualitative relative strengths are, from highest to lowest, J23100, J23106, and J23103^5^. For RBSs, several different types were chosen, including bicistronic and ribozyme-insulated designs. The standard RBSs had qualitative relative strengths ordered B0034m, B0032m, and B0033m. The bicistronic and insulated RBSs BCD2, RiboJ+BCD2, and RiboJ+UTR1 were expected to be high expression, but with unknown relative magnitude.

To assess expression strength in *E. coli* and the other three hosts, we built a library of fluorescent reporter strains under the control each combination of the three promoters and six RBSs. Although this set is small, we developed a liquid handling protocol for assembly that could be scaled to the size of the CIDAR parts set (Methods). We then transformed this set of 18 expression units into each host, quantified fluorescence intensity using flow cytometry, calibrated fluorescence intensity to an external standard (molecules of equivalent fluorescein, or MEFL), and subtracted cell autofluorescence. The expression strengths for a portion of this combinatorial library have previously been measured in *E. coli* in the original CIDAR vectors (Figure 2B, values extrapolated from the CIDAR MoClo supplementary information^5^). When compared to the openCIDAR expression values for *E. coli* (Figure 2C), we found that the relative expression strengths for combinations of parts characterized in CIDAR and openCIDAR vectors were consistent. In general, we found that expression strengths were qualitatively similar across the other hosts *P. putida* KT2440 (Figure 2D), *C. necator* H16 (Figure 2E), and *K. nataicola* DS12 (Figure 2F). This was true despite the copy number variation across hosts. However, the violin plots revealed low-expressing sub-populations with fluorescence intensity close to the empty vector control in *P. putida*, while this phenomenon was not observed in *E. coli*.

To better assess part performance across hosts, we first plotted all expression values on the same chart, which is possible because all values are expressed in terms of MEFL (Figure 3A). Unexpectedly, global expression values for all constructs trended higher in *C. necator* and *P. putida* than in *E. coli*. Then, Pearson correlation coefficients were calculated for the constructs across hosts (Figure 3B). Expression values for the same constructs in *E. coli* and *P. putida* were well-correlated (Pearson coefficient R= 0.84). Expression levels were well-correlated between *C. necator* and *E. coli* (R=0.83). In contrast, expression levels were poorly correlated between *K. nataicola* and *E. coli* (R=0.33). This demonstrates that although a range of expression values are possible in *K. nataicola*, the same combinations of promoters and RBS cannot be used to achieve similar expression levels. This necessitates an MEFL lookup table (LUT), which can be used to transfer a genetic design from one organism to another (Figure 4). With a LUT, equivalent parts in a new organism can be matched to known part sets, or part sets which achieve a desired expression level can be selected. This is made possible by the MEFL standard expression unit.

**Figure 3:**
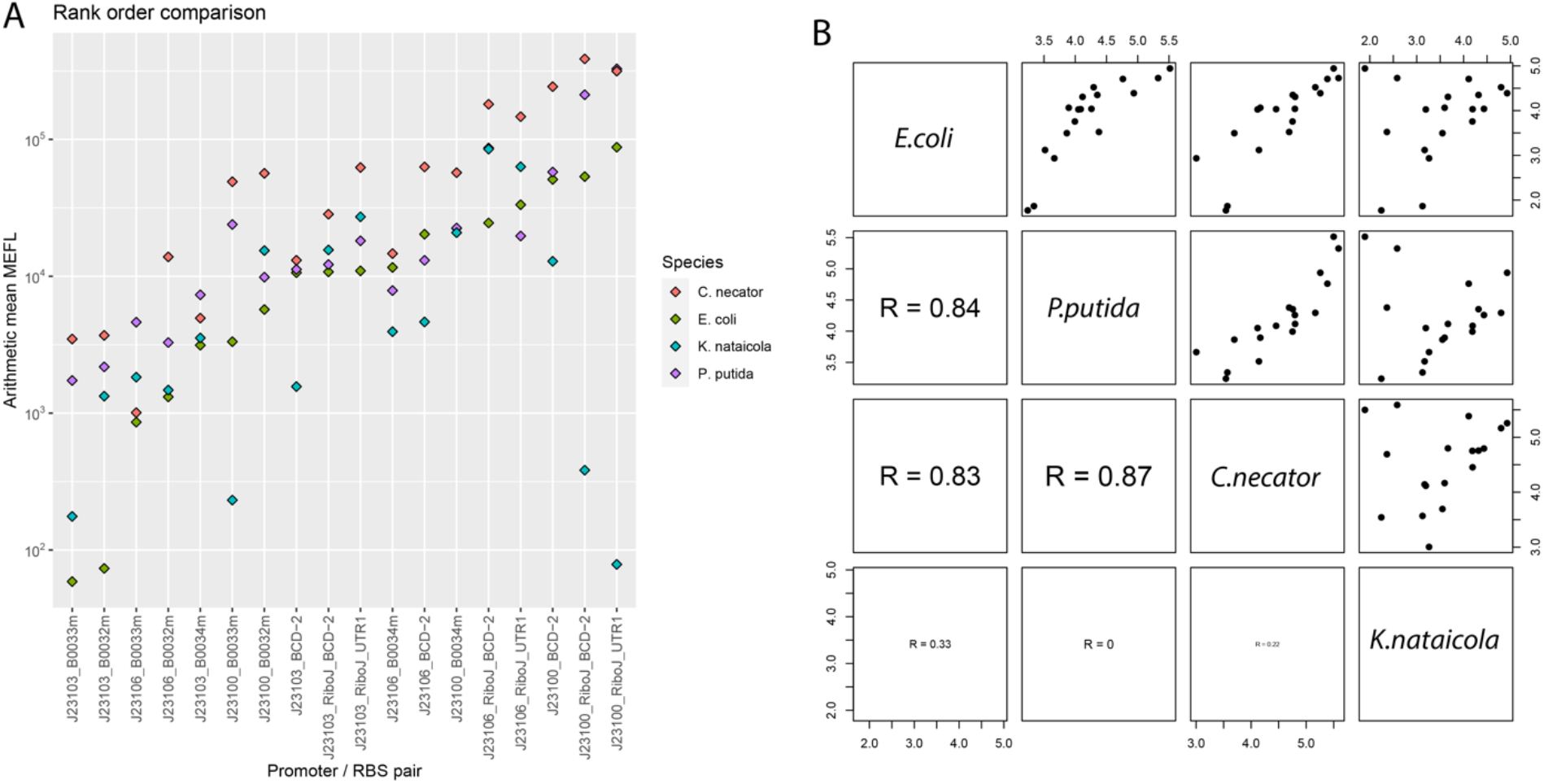
Comparing and correlating part function across hosts. **A**. Rank order comparison of MEFL for each construct in each host. This shows a rough global correlation for expression in all species. Divergent values are largely found only in *K*.*nataicola*. **B**. Species-species correlations. The species is in the diagonal, with the MEFL values in each species plotted in the top right boxes and the correlation coefficients in the bottom left boxes.

**Figure 4:**
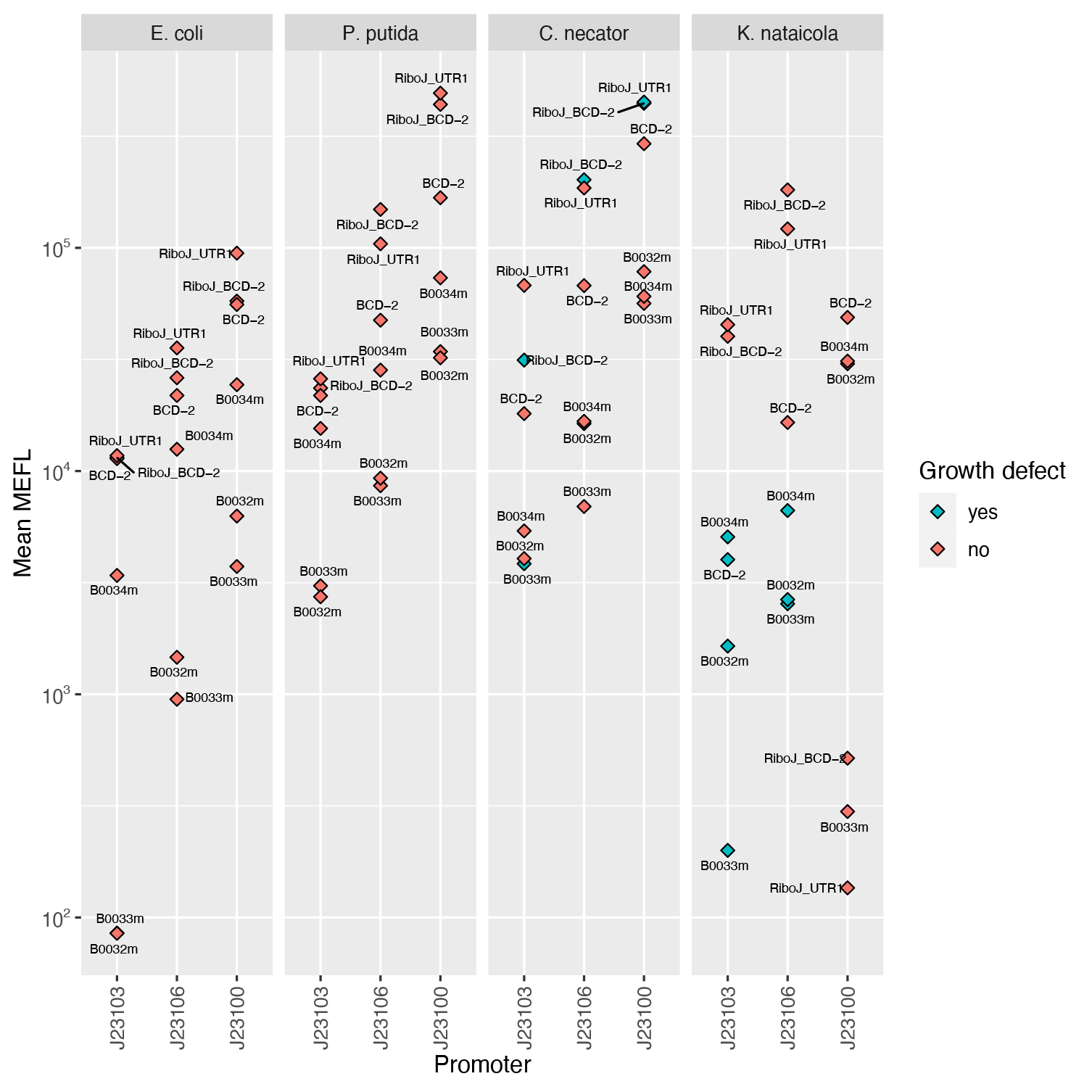
MEFL lookup table (LUT) for openCIDAR. By matching MEFL across organisms, researchers may design genetic constructs that achieve the same expression values across different organisms.

**Figure 5:**
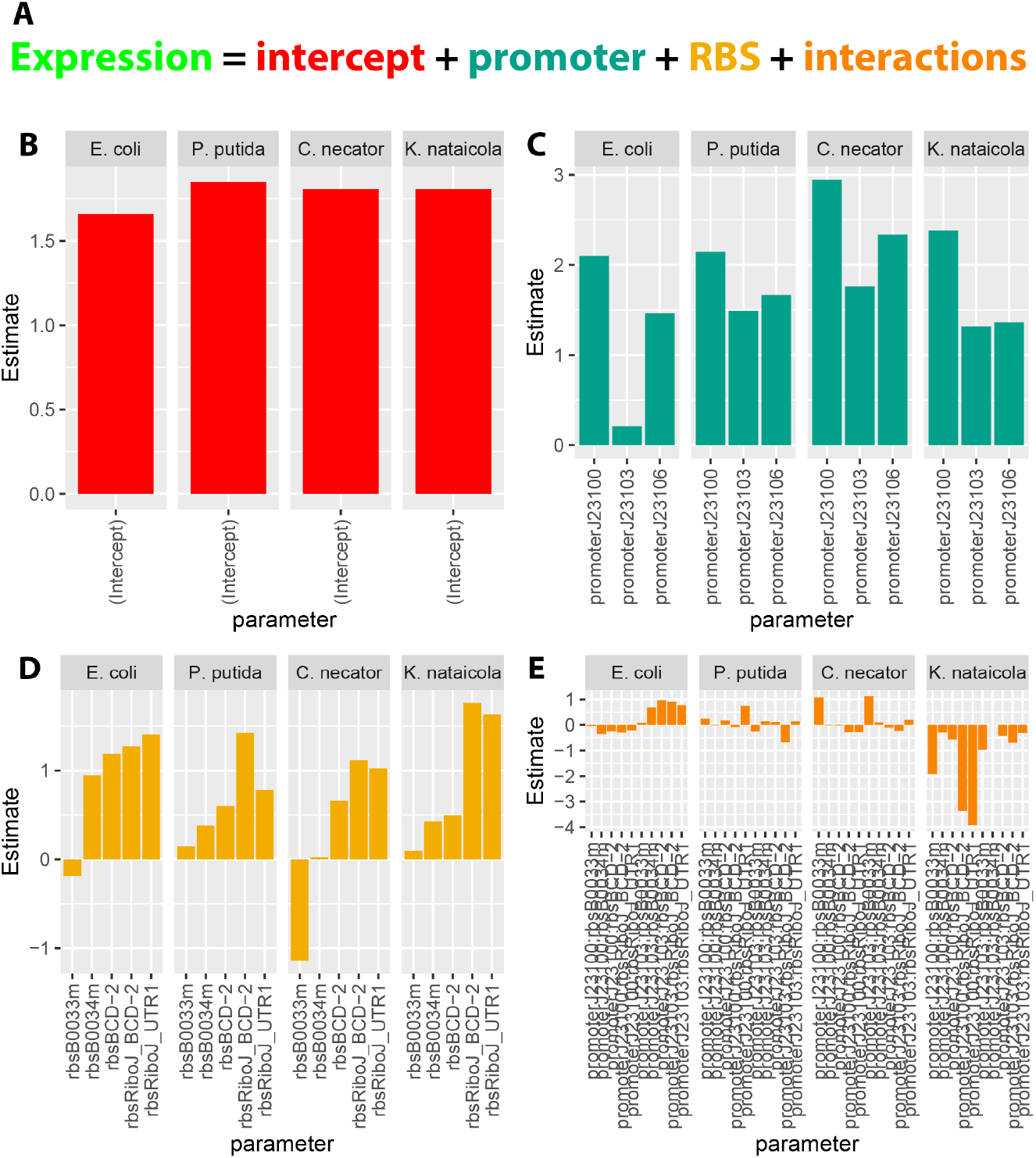
Linear models for part performance across species. **A**. General form of the linear models. Expression is the sum of an intercept term plus estimates computed for each promoter, RBS, and the interactions between them. Due to high correlation between variables (i.e. very similar expression between parts), some parameters were not estimated (see Methods). **B**. Bar chart of the intercept values, interpreted as background fluorescence, were similar between species. **C**. Bar chart of the promoter coefficients. The contribution of each promoter to expression had conserved rank order in each species, however the magnitudes varied. Notably, the expected very low expression of the J23103 promoter in *E. coli* was not conserved. **D**. Bar chart of the RBS coefficients. Variation in the contribution of the RBS to expression between species appears greater than the variation in the contribution of promoters. **E**. Bar chart of the interaction terms between promoters and RBSs. Interactions between promoter and RBS terms were very high in *K. nataicola*, especially for those terms including the J23100 promoter.

Truly interchangeable parts should have negligible interactions with other elements. To investigate interactions, linear regression models of GFP expression by promoter, RBS, and the interactions between the two (all other construct elements were constant) were fit for each organism. The coefficients of each term (log_10_ transformed) were compared as a proxy for understanding the relative effect size of each term. The individual effect sizes of the intercept (baseline fluorescence), promoters, and RBSs were usually the dominant factors, indicating independent contributions to the total MEFL. For *K. nataicola*, however, the interaction between promoter J23100 and several RBSs had coefficients equivalent or larger than the promoter coefficients themselves. We conclude that promoter J23100 is not “well-behaved” in *K. nataicola*, that is it does not function the same across hosts, even though it still produces functional transcription units. This is a notable finding as promoter J23100 is a very commonly used strong promoter, and is likely to be one of the first choices for synthetic biologists working in novel *Pseudomonadota*. This interaction would be difficult to detect without a combinatorial approach and quantitative analysis. The behavior of this single promoter is largely the cause of the lack of correlation between *K. nataicola* and the other organisms in Figure 3B.

## DISCUSSION

As new organisms are identified as chassis of interest for synthetic biology, appropriate tools and part sets for controlling gene expression need to be identified. Here, we demonstrate a method for rapidly identifying appropriate expression parts for multiple organisms from a single parts set. This obviates the need to develop new part sets by genome mining or other methods. This should enable researchers to more rapidly onboard new organisms.

Previous studies have explored the range of fluorescent protein expression achievable in nonconventional prokaryotes by varying promoters with a consistent RBS, or RBS with a consistent promoter^2,4,7,27^. Few, if any, studies have explored fully combinatorial sets of both promoters and RBSs across several hosts from diverse lineages. This, coupled with the standard MEFL measurement, enabled analyses not previously possible, with several interesting outcomes.

First, we noted that expression of the same DNA across different hosts resulted in MEFL values that were generally well-correlated. However, *P. putida, C. necator*, and *K. nataicola* all had a higher average expression strength than *E. coli* which did not correlate with copy number. This suggests that other host factors besides plasmid copy number may be contributing to expression. Furthermore, these results show that at least in some bacteria, it may be more difficult to achieve low expression rather than high expression.

Second, we observed that the promoters are largely functional across hosts. This is not unique to bacteria – this has been observed in yeast^25^ and animals^26^. Yet, the yeast study clearly showed an evolutionary distance trend where sufficiently distant promoters were no longer functional (promoters from *Yarrowia lipolytica* and *Schizosaccharomyces pombe* were not functional in *Saccharomyces cerevisiae*). In contrast, we observed a breakdown in expression patterns but not a nonfunctional barrier, even for distantly related bacteria. The lack of a nonfunctional barrier could be due to the efficiency and simplicity of prokaryotic gene expression, the parts being synthetic rather than host derived, greater conservation of prokaryotic gene expression machinery, or some other mechanism. Differences in expression levels are possibly due to differences in transcription factor binding affinity, ribosome binding affinity, and global protein expression control within the cell.

To enable similar function of genetic designs across hosts, we propose designs should be transferred using a lookup table that matches equivalent expression parts across hosts. This would account for the variability of the same DNA across hosts. Rebuilding a design for a new host is particularly facile in the openCIDAR system compared to other systems due to the modular cloning framework and availability of a large parts set, although the characterization strategy could be used to translate between different part collections as well, since the comparison is made with MEFL, not arbitrary units. Furthermore, by using combinatorial experiments and linear regression, it was possible to identify the divergent behavior of the J23100 promoter in *K. nataicola*. This demonstrates that it is possible to debug parts and derive design rules in each organism, in addition to the lookup table, that can inform future designs.

While the parts function across the major classes of *Pseudomonadota*, it is important to note that several factors such as transposons, host restriction systems, and growth rate could limit the application of this parts set in a potential new host. We know this is at least true of *Vibrio natriegens* (communication with Tanya Tschirhart), as pBBR1 is particularly unstable even though functional^27^. However, the lookup table approach would still be useful to translate *E. coli* designs to a *V. natriegens* expression system, as long as all parts are measured in MEFL. Furthermore, others are developing approaches to overcome host limitations like tools that evade restriction-modification systems^29^. Despite these limitations, openCIDAR should at least work well in all species that can stably maintain pBBR1.

In conclusion, we have defined a system that enables transfer of accessible, well-characterized parts to new chassis. By screening a modest library, we achieved constitutive expression over three orders of magnitude in each species studied. Coupled with recent advances in scalable and high-throughput electroporation devices^28^, our approach can make automated, high-throughput, modular DNA assembly in many nonconventional prokaryotes an achievable and practical workflow. This is because only one part set is required for a large group of related bacteria – hinting that only a few parts sets may be needed to enable expression across the tree of life.

## MATERIAL AND METHODS

### Phylogenetic placement of pBBR1-compatible bacteria

A literature search was conducted to identify bacterial species reported to be transformable with the pBBR1 plasmid and derivatives. These species are reported in **Error! Reference source not found**.. Representative genomes for each species were accessed from the NCBI RefSeq database, and a species tree showing phylogenetic relationships was constructed from genomic protein sequences using the OrthoFinder tool^30,31^.

### Strains and media

All cloning was performed in NEB 5-alpha Competent *Escherichia coli* (New England BioLabs C2987U). The *Pseudomonas putida* KT2440 strain was purchased from ATCC (#47054). The *Cupriavidus necator* H16 strain was purchased from ATCC (#17699). The *Komagataeibacter nataicola* DS12 strain was isolated from a kombucha tea culture (Urban Farm, Portland ME) by Elizabeth van Zyl of the Coburn lab and provided as a gift. Routine culture of *E*.*coli* for cloning purposes was performed in Miller’s LB Broth (Fisher BP1426-2) in 14mL Falcon tubes (VWR 60819-761) at 37°C on a rotating drum. Routine culture of *P*.*putida* and *C*.*necator* was performed in Miller’s LB Broth using 50mL conical tubes in an orbital incubator shaker at 30°C, 220 RPM. Routine culture of *K*.*nataicola* was performed in Hestrin-Schramm media^32^ (5 g/L Bacto Peptone(BD 211677), 5 g/L Difco Yeast Extract (BD 210929), 1.15 g/L citric acid (Sigma-Aldrich C0759), 2.7 g/L sodium phosphate dibasic anhydrous (Sigma-Aldrich S5136), 20 g/L glucose (Sigma-Aldrich G7021)) using 125mL shake flasks in an orbital incubator shaker at 30°C, 220 RPM. Solid media plates were prepared using the previously described media supplemented with 20g/L (*E*.*coli, P*.*putida, C*.*necator*) or 15 g/L (*K*.*nataicola*) agar (Sunrise Science Products 1910-1KG). For antibiotic selection and plasmid maintenance, kanamycin (Alfa Aesar J61272) was added at 50 μg/mL (*E*.*coli, P*.*putida*), 100 μg/mL (*K*.*nataicola*) and 200 μg/mL (*C*.*necator*). For identification of background colonies in cloning, X-Gal was supplemented to the selection media (Thermo Scientific R0941).

The CIDAR MoClo Parts Kit was a gift from Douglas Densmore (Addgene kit # 1000000059). CIDAR MoClo Extension, Volume I was a gift from Richard Murray (Addgene kit #1000000161). pSEVA331Bb was a gift from Tom Ellis (Addgene plasmid # 78269 ; http://n2t.net/addgene:78269 ; RRID:Addgene_78269).

### Cloning of new destination vector series

To construct the Level 1 series, the BBR1 origin of replication was amplified from pSEVA331Bb (Addgene #78269). A silent mutation was introduced in the Rep protein to remove a BbsI restriction site (G186A). The kanamycin resistance cassette was amplified from pAN861. The CIDAR MoClo cloning sites were amplified from DVK_AE, DVK_EF, DVK_FG and DVK_GH. The fragments containing the origin of replication, kanamycin resistance gene and cloning site, along with appropriate flanking sequences, were combined by Gibson Assembly (NEB E2621S) to form pBBR1_DVK_AE, pBBR1_DVK_EF, pBBR1_DVK_FG and pBBR1_DVK_GH. To construct the Level 2 series, the fragment containing the BBR1 origin of replication was combined with a chloramphenicol resistance cassette from pY122 or a spectinomycin resistance cassette from pICH41308 and the cloning site from DVA_AF, DVA_AG and DVA_AH to form pBBR1_DVS_AF, pBBR1_DVS_AG, pBBR1_DVS_AH, pBBR1_DVC_AF, pBBR1_DVC_AG and pBBR1_DVC_AH. Correct assembly was confirmed by Sanger sequencing. All new destination vectors constructed for this study are summarized in **Error! Reference source not found**..

### Automated cloning pipeline

The CIDAR MoClo Parts Kit (#1000000059) and CIDAR MoClo Extension, Volume I (#1000000161) were purchased from Addgene. Plasmid DNA was purified from *E*.*coli* host strains using the QIAprep Spin Miniprep kit (Qiagen 27106). Plasmid DNA was quantitated using a NanoDrop spectrophotometer and diluted to 20 fmol/μL in nuclease-free water (VWR 02-0201-0500). An appropriate volume of reaction mastermix was prepared with the following composition: 20% v/v BsaI (New England BioLabs R3733S) or BbsI (Thermo Scientific ER1012), 8% v/v T4 DNA ligase HC (Promega M1794), 20% v/v T4 DNA ligase buffer (Promega), 52% v/v nuclease-free water. This mastermix was transferred to a Labcyte Echo 384 LDV plate (001-13070), along with an appropriate volume of each DNA part. For a five part assembly, 200nL of each part (4 fmol) was transferred to an Eppendorf twin.tec 96-well PCR plate (951020427), along with 1μL of mastermix using a Labcyte Echo 525 acoustic liquid handler. The transfers were programmed using the Echo Plate Reformat software. After completion of transfer, the PCR plate was sealed with PCR film (VWR 60941-078) and the following reaction was carried out on a thermocycler: 37°C for 90 seconds then 16°C for three minutes cycled 25 times, followed by five minutes and 50°C and 10 minutes at 80°C. The plate was then transferred to an Eppendorf epMotion 5075t liquid handler, where the following operations were performed: 10μL NEB 5-alpha Competent *E. coli* (C2987U) cells added to each well, shaken at 500 RPM for 15 seconds, incubated for 15 minutes in a block pre-frozen at -20°C, heat shocked at 42°C for 30 seconds, moved back to cold block for 5 minutes, 100μL SOC added, incubated at 37°C for one hour. Subsequently, 10μL of transformation mixture was transferred to unused wells of the original Echo 384 well plate, and that plate was transferred to the Echo 525 acoustic liquid handler. Transformed cells were transferred to pre-warmed (37°C) antibiotic selection media plates (OmniTray, Thermo Scientific 264728) in a pseudo-96 well pattern, with one 25nL droplet per spot and 18 spots per pseudo-well. Plates were incubated overnight at 37°C and white colonies were picked. Positive colonies were confirmed for correct assembly by enzymatic digestion (with BbsI for level 1 assemblies) followed by gel electrophoresis band size analysis (Invitrogen G401001).

### Transformation of Komagataeibacter nataicola

The method for transformation of *K*.*nataicola* was adapted from Florea et al^22^. A source plate of *K*.*nataicola* was streaked out on HS media, incubated for 3-4 days at 30°C and then stored at RT for up to a month. A whole colony was picked from this plate and inoculated into a preculture of 10mL HS media with 0.2% v/v cellulase (Sigma Aldrich C2730) in a 125mL flask. When grown on solid media, *K*.*nataicola* forms hard, connected networks, making the difference between a “whole colony” and a “small chunk” somewhat subjective. The preculture was incubated with shaking at 30°C, 220 RPM for 2-3 days until quite cloudy (OD_600_∼1-2). From this preculture, two 50mL cultures (in 250mL flasks) were inoculated at OD=0.005 and 0.010. These flasks were incubated with shaking at 30°C, 220 RPM overnight. The next day, the flask in the range of OD=0.4-0.7 at the time of transformation was used. Prior to transformation, a swinging-bucket centrifuge was pre-chilled to 4°C for at least 30 minutes, until the centrifuge cups were cold to the touch. Solutions of 1mM HEPES (diluted from Fisher BP299) and 15% glycerol (Fisher BP229) were stored at 4°C and transferred to ice buckets as the centrifuge was chilled. Cells were decanted into a 50mL conical tube and transferred to ice for 10 minutes. From this step forward, all procedures were performed on ice. Cells were pelleted at 3000xg for 5 minutes, and the supernatant was decanted and discarded. The pellet was washed twice in 10mL HEPES with the following method: gently resuspend the pellet by tapping, resuspend in 1mL using a P1000 (breaking up any clumps), resuspend in the remaining 9mL using a 10mL serological pipette, then pellet at 3000xg for 5 minutes. During this these washes, 1mm electroporation cuvettes (Molecular BioProducts 5510) were stored on ice and labelled, along with 0.2 mL PCR tubes (USA Scientific 1402-4700) for each transformation. Plasmid DNA (up to 200 ng, maximum volume 4μL) was transferred to PCR tubes. The final washed pellet was resuspended in 1mL ice-cold 15% glycerol, and 100μL competent cell suspension was added to the DNA in the PCR tubes. Any remaining competent cells were aliquoted into PCR strips and stored at -80°C for later use. The tubes were gently tapped to mix and briefly centrifuged to collect in the tube bottom, and then incubated on ice for 10 minutes. Each mixture was then transferred to the corresponding electroporation cuvette, which was dried and electroporated at 2.5 kV, 400 Ohm resistance, 25 μF capacitance. Recovery media (1mL HS with 0.2% cellulase) was immediately added to the cuvette. Following all transformations, cells in recovery media were transferred to 14mL round bottom Falcon tubes and incubated with shaking at 30°C, 220 RPM overnight. Transformation efficiency varied from construct to construct. Typically, 100μL of recovered cells were spread on a 10cm dish containing selection media and incubated at 30°C for 3-4 days. If this procedure did not yield colonies, the transformation was repeated and the entire volume of transformed cells was centrifuged and resuspended in 200μL HS with the appropriate antibiotic, then plated on selective media. Colonies were picked into 10mL HS with 0.2% cellulase and the appropriate antibiotic, grown at 30°C, 220 RPM until cloudy (typically 1-2 days but up to 4 for slow-growing strains), and preserved as glycerol stocks (500μL cell culture plus 500μL 50% glycerol for final glycerol concentration of 25%).

### Transformation of Pseudomonas putida

The method for transformation of *P*.*putida* was adapted from the method described by Elmore et al^33^. A single colony was inoculated into 10mL of LB in a 50mL conical tube and incubated with agitation at 30°C, 220 RPM overnight. The protocol was scaled up as needed by adding more tubes of 10mL culture. The cell culture was pelleted at 3,200xg for 10 minutes, and washed 3x with 5mL of 10% v/v glycerol (room temperature). This pellet was resuspended in 200μL of 10% glycerol per 10mL starting culture. Aliquots of 50μL in PCR strips were stored at -80°C. At the time of transformation, aliquots were thawed at RT for 5 minutes, and 10μL of these competent cells were diluted into 40μL of 10% glycerol in PCR strips. Plasmid DNA was added to these cells (50-150ng in 1-2μL). Cells and DNA were mixed by flicking the tubes, which were then briefly spun down and incubated at RT for 5 minutes. The mixture was transferred to a 1mm electroporation cuvette and electroporated at 1600V. The time constant was typically 5-6ms. SOC recovery media (250μL) was immediately added to the cuvette following electroporation. The recovering cells were transferred to a 14mL round bottom Falcon tube and incubated at 30°C, 220 RPM for 90 minutes. Following recovery, 5μL of transformation mixture was added to a selective media plate and streaked out for single colonies. Plates were incubated overnight at 30°C and single colonies were picked.

### Transformation of Cupriavidus necator

The method for transformation of *C*.*necator* was adapted from the method described by Peabody et al. in an unpublished manuscript shared with our lab by Adam Guss. The culture was agitated at 220 RPM rather than 250. In addition, the DNA was incubated with the cells for 5 minutes at RT. Post-transformation recovery was carried out in a 14mL round bottom Falcon tube rather than a microcentrifuge tube. All other aspects of the transformation protocol were carried out as described. With the plasmid constructs described here, 100ng DNA was used for each transformation.

### Flow cytometry

The method for cell characterization by flow cytometry was adapted from Andrews et al^34^. For *E*.*coli* and *P*.*putida*, single colonies were picked into wells of a 96-well V-bottom plate (Thermo Fisher 249662) containing 100μL LB with the appropriate antibiotic. This plate was incubated overnight with shaking at 1000 RPM, 37°C for *E*.*coli*, 30°C for *P*.*putida*. Cells were then diluted into fresh media (2μL into 98μL) in triplicate and grown for 6 hours. From this culture, 10μL was transferred into a plate containing 190μL diH_2_O with 2mg/mL kanamycin. This plate was incubated at room temperature for 1 hour and used directly for flow cytometry.

For *C*.*necator* and *K*.*nataicola*, single colonies were picked into 5mL LB or HS (respectively) in a 50mL conical tube and incubated with shaking at 30°C, 220 RPM until quite cloudy. Cells were then diluted into 5mL media in triplicate (5μL for *C*.*necator*, 25 μL *K*.*nataicola*) and incubated until slightly cloudy. From these culture, 10μL was transferred into a plate containing 190μL diH_2_O with 2mg/mL kanamycin. This plate was incubated at room temperature for 1 hour and used directly for flow cytometry.

A Beckman Coulter CytoFLEX S flow cytometer was used with the following parameters: GFP channel configured using blue laser (488nm, 50mW) with 525/40nm band pass filter, gain 90. Gating thresholds FSC 1500 (gain 68) and SSC 1000 (gain 56). SpheroTech Rainbow Calibration particles (URCP-38-2K) and destination vector-only (pBBR1_DVK_AE) control were used for calibration and subsequent analysis. Fluorescent intensities were converted to MEFL using the FlowCal toolkit in a custom Python script^35^.

### Plasmid copy number determination by ddPCR

Cell cultures were grown to mid-exponential phase and frozen at -80°C prior to analysis. Sample preparation was adapted from Jahn et al^36^. Cell samples were thawed on ice and diluted into 1mL nuclease-free water at 1:500 or 1:250 dilutions. From this diluted sample, 44μL were transferred to PCR strips and boiled at 95°C for 20 minutes for lysis. BsaI (1 μL) and 10x CutSmart buffer (5 μL) were added to each sample, and the plasmid and genomic DNA was digested at 37°C for 30 minutes. Reaction mixtures were prepared by combining the following reagents per wells to BioRad ddPCR plates: 5.5 μL 4x ddPCR SuperMix; 4.4 μL lysed, digested sample; 1.1 μL each probe (5 μM stock) and primer (18 μM stock); and 5.5 μL nuclease-free water. The plate was heat-sealed in foil, vortexed vigorously and centrifuged briefly. Droplets were generated using the BioRad AutoDropletGenerator using the manufacturer’s recommended reagents and consumables. Droplets were read using a QX200 droplet reader with the QuantaSoft software and analyzed using the QX Manager Standard software.

Primers and probes were designed for single-copy genomic targets of each strain as well as the BBR1 *rep* protein. Genomic targets were labelled with the FAM fluorophore and BBR1 plasmid targets were labelled with the HEX fluorophore for multiplexing. The *E*.*coli* cysG, and *P*.*putida* ileS primer/probe sets were used as described by Jahn *et al*^37^. Primers and probes for the mutated BBR1 *rep* protein, *C*.*necator* panC target^6^ and the *K*.*nataicola* cysG target were designed using NCBI Primer-BLAST. Primer details are included in **Error! Reference source not found**..

## RESEARCH DATA AVAILABILITY

Digital data files related to this study are available from GitHub repository github.com/kevinwkeating/openCIDAR. Data available in this repository includes plasmid sequences, flow cytometry raw data and analysis scripts, and ddPCR analysis exports. Destination vectors and fluorescent reporter cassettes have been deposited with Addgene (**Error! Reference source not found**.).

## ABBREVIATIONS

MEFL: molecules of equivalent fluorescein
ddPCR: droplet digital polymerase chain reaction

## ACKNOWLEDGEMENTS

Violet Smiarowski and Hannah Peloquin assisted with molecular biology tasks. Nisreen Aljumaili assisted with development of the automated cloning pipeline. The authors thank Tanya Tschirhart for initial experiments and personal communication regarding *V. natriegens*.

## REFERENCES

1. Blombach, B., Grünberger, A., Centler, F., Wierckx, N. & Schmid, J. Exploiting unconventional prokaryotic hosts for industrial biotechnology. Trends Biotechnol. 40, 385–397 (2022).

2. Goosens, V. J. et al. Komagataeibacter Tool Kit (KTK): A Modular Cloning System for Multigene Constructs and Programmed Protein Secretion from Cellulose Producing Bacteria. ACS Synth. Biol. 10, 3422–3434 (2021).

3. Moore, S. J. et al. EcoFlex: A Multifunctional MoClo Kit for E. coli Synthetic Biology. ACS Synth. Biol. 5, 1059–1069 (2016).

4. Teh, M. Y. et al. An Expanded Synthetic Biology Toolkit for Gene Expression Control in Acetobacteraceae. ACS Synth. Biol. 8, 708–723 (2019).

5. Iverson, S. V., Haddock, T. L., Beal, J. & Densmore, D. M. CIDAR MoClo: Improved MoClo Assembly Standard and New E. coli Part Library Enable Rapid Combinatorial Design for Synthetic and Traditional Biology. ACS Synth. Biol. 5, 99–103 (2016).

6. Ehsaan, M., Baker, J., Kovács, K., Malys, N. & Minton, N. P. The pMTL70000 modular, plasmid vector series for strain engineering in Cupriavidus necator H16. J. Microbiol. Methods 189, 106323 (2021).

7. Johnson, A. O., Gonzalez-Villanueva, M., Tee, K. L. & Wong, T. S. An Engineered Constitutive Promoter Set with Broad Activity Range for Cupriavidus necator H16. ACS Synth. Biol. 7, 1918–1928 (2018).

8. Alagesan, S. et al. Functional Genetic Elements for Controlling Gene Expression in Cupriavidus necator H16. Appl. Environ. Microbiol. 84, e00878–18 (2018).

9. Silva-Rocha, R. et al. The Standard European Vector Architecture (SEVA): a coherent platform for the analysis and deployment of complex prokaryotic phenotypes. Nucleic Acids Res. 41, D666–D675 (2013).

10. García-Gutiérrez, C. et al. Multifunctional SEVA shuttle vectors for actinomycetes and Gram-negative bacteria. MicrobiologyOpen 9, e1024 (2020).

11. Arnold, B. J., Huang, I.-T. & Hanage, W. P. Horizontal gene transfer and adaptive evolution in bacteria. Nat. Rev. Microbiol. 20, 206–218 (2022).

12. Choi, K.-H. & Kim, K.-J. Applications of Transposon-Based Gene Delivery System in Bacteria. J. Microbiol. Biotechnol. 19, 217–228 (2009).

13. Kushwaha, M. & Salis, H. M. A portable expression resource for engineering cross-species genetic circuits and pathways. Nat. Commun. 6, 7832 (2015).

14. Brophy, J. A. N. et al. Engineered integrative and conjugative elements for efficient and inducible DNA transfer to undomesticated bacteria. Nat. Microbiol. 3, 1043–1053 (2018).

15. Mutalik, V. K. et al. Precise and reliable gene expression via standard transcription and translation initiation elements. Nat. Methods 10, 354–360 (2013).

16. Cook, T. B. et al. Genetic tools for reliable gene expression and recombineering in Pseudomonas putida. J. Ind. Microbiol. Biotechnol. 45, 517–527 (2018).

17. Nikel, P. I. & de Lorenzo, V. Pseudomonas putida as a functional chassis for industrial biocatalysis: From native biochemistry to trans-metabolism. Metab. Eng. 50, 142–155 (2018).

18. Panich, J., Fong, B. & Singer, S. W. Metabolic Engineering of Cupriavidus necator H16 for Sustainable Biofuels from CO2. Trends Biotechnol. 39, 412–424 (2021).

19. Zhang, H. et al. Complete genome sequence of the cellulose-producing strain Komagataeibacter nataicola RZS01. Sci. Rep. 7, 4431 (2017).

20. Beal, J. et al. TASBE Flow Analytics: A Package for Calibrated Flow Cytometry Analysis. ACS Synth. Biol. 8, 1524–1529 (2019).

21. Beal, J. et al. Quantification of bacterial fluorescence using independent calibrants. PLOS ONE 13, e0199432 (2018).

22. Florea, M. et al. Engineering control of bacterial cellulose production using a genetic toolkit and a new cellulose-producing strain. Proc. Natl. Acad. Sci. 113, E3431–E3440 (2016).

23. Kiewiet, R., Kok, J., Seegers, J. F. M. L., Venema, G. & Bron, S. The Mode of Replication Is a Major Factor in Segregational Plasmid Instability in Lactococcus lactis. Appl. Environ. Microbiol. 59, 358– 364 (1993).

24. Tee, K. L. et al. An Efficient Transformation Method for the Bioplastic-Producing “Knallgas” Bacterium Ralstonia eutropha H16. Biotechnol. J. 12, 1700081 (2017).

25. Zeevi, D. et al. Molecular dissection of the genetic mechanisms that underlie expression conservation in orthologous yeast ribosomal promoters. Genome Res. 24, 1991–1999 (2014).

26. Fisher, S., Grice, E. A., Vinton, R. M., Bessling, S. L. & McCallion, A. S. Conservation of RET Regulatory Function from Human to Zebrafish Without Sequence Similarity. Science 312, 276–279 (2006).

27. Tschirhart, T. et al. Synthetic Biology Tools for the Fast-Growing Marine Bacterium Vibrio natriegens. ACS Synth. Biol. 8, 2069–2079 (2019).

28. Huang, P.-H. et al. M-TUBE enables large-volume bacterial gene delivery using a high-throughput microfluidic electroporation platform. PLOS Biol. 20, e3001727 (2022).

29. Riley, L. A., Ji, L., Schmitz, R. J., Westpheling, J. & Guss, A. M. Rational development of transformation in Clostridium thermocellum ATCC 27405 via complete methylome analysis and evasion of native restriction–modification systems. J. Ind. Microbiol. Biotechnol. 46, 1435–1443 (2019).

30. Emms, D. M. & Kelly, S. OrthoFinder: solving fundamental biases in whole genome comparisons dramatically improves orthogroup inference accuracy. Genome Biol. 16, 157 (2015).

31. Emms, D. M. & Kelly, S. OrthoFinder: phylogenetic orthology inference for comparative genomics. Genome Biol. 20, 238 (2019).

32. Schramm, M. & Hestrin, S. Synthesis of cellulose by Acetobacter xylinum. 1. Micromethod for the determination of celluloses*. Biochem. J. 56, 163–166 (1954).

33. Elmore, J. R. et al. Production of itaconic acid from alkali pretreated lignin by dynamic two stage bioconversion. Nat. Commun. 12, 2261 (2021).

34. Woodruff, L. B. A. et al. Registry in a tube: multiplexed pools of retrievable parts for genetic design space exploration. Nucleic Acids Res. 45, 1553–1565 (2017).

35. Castillo-Hair, S. M. et al. FlowCal: A User-Friendly, Open Source Software Tool for Automatically Converting Flow Cytometry Data from Arbitrary to Calibrated Units. ACS Synth. Biol. 5, 774–780 (2016).

36. Jahn, M. et al. Accurate Determination of Plasmid Copy Number of Flow-Sorted Cells using Droplet Digital PCR. Anal. Chem. 86, 5969–5976 (2014).

37. Jahn, M., Vorpahl, C., Hübschmann, T., Harms, H. & Müller, S. Copy number variability of expression plasmids determined by cell sorting and Droplet Digital PCR. Microb. Cell Factories 15, 211 (2016).

