## Supplemental tables and figures for "Systematic part transfer by extending a modular toolkit to diverse bacteria"

### SUPPLEMENTARY FIGURES

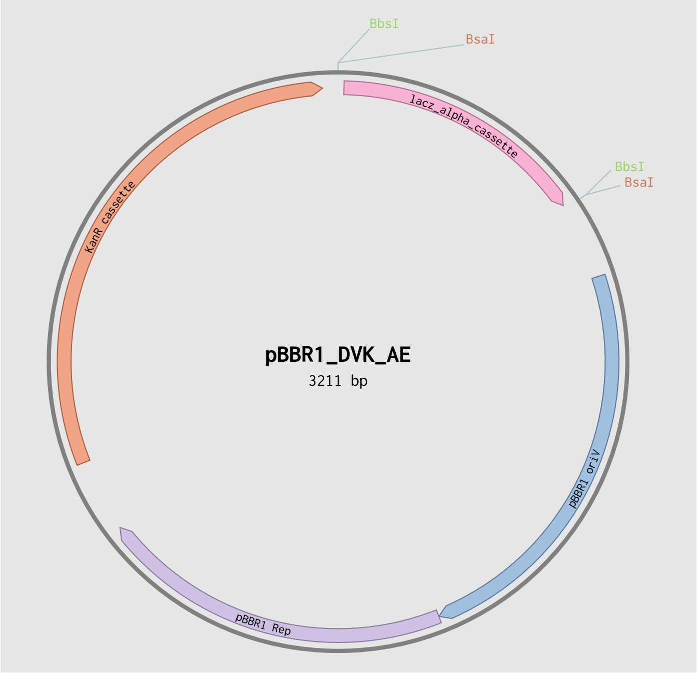

**Supplementary Figure 1: Plasmid map of pBBR1_DVK_AE**

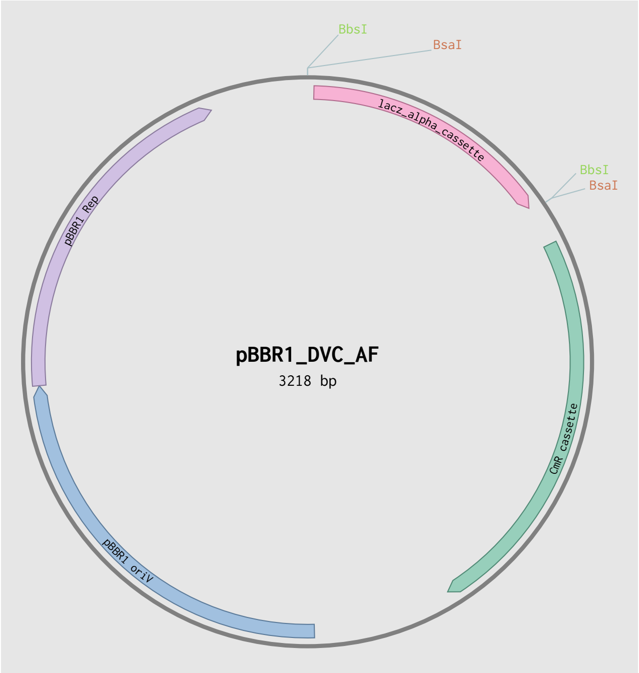

**Supplementary Figure 2: Plasmid map of pBBR1_DVC_AF**

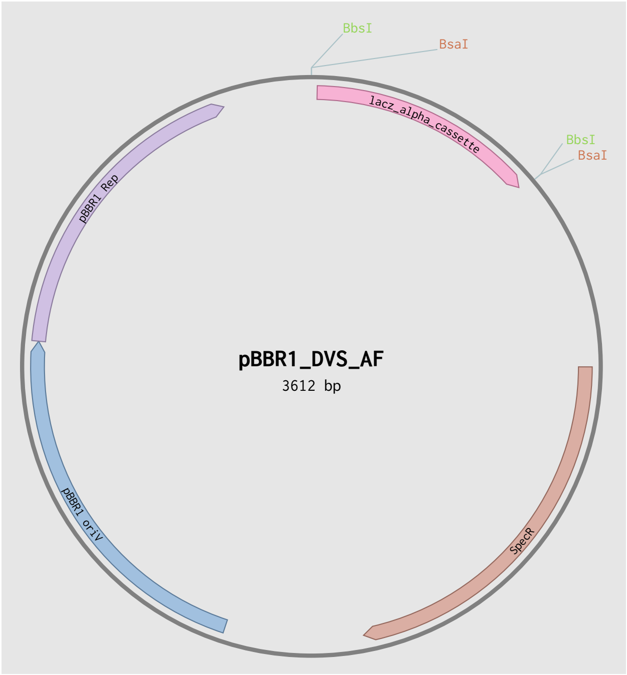

**Supplementary Figure 3: Plasmid map of pBBR1_DVS_AF**

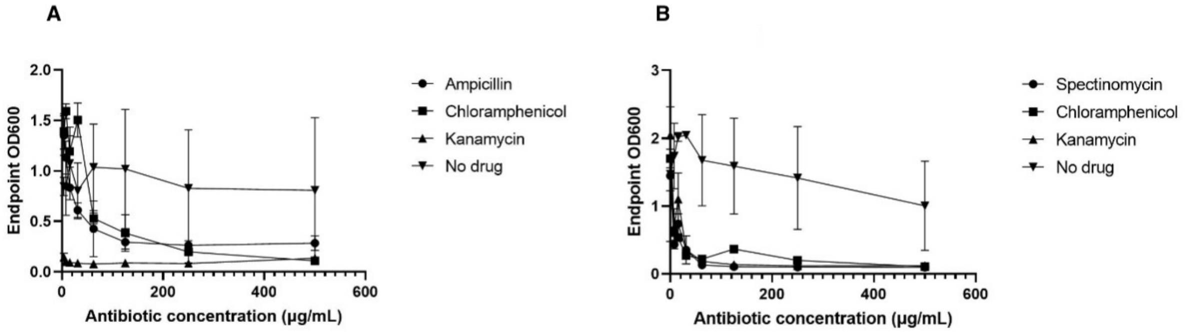

**Supplementary Figure 4: *K.nataicola* DS12 antibiotic sensitivity**

A: Cells from an overnight culture of *K.nataicola* DS12 were inoculated into HS media with cellulase and the given concentration of antibiotic. Cells were grown to stationary phase (no drug control) and the optical density was read. Kanamycin inhibited growth, even at the lowest tested concentrations. Chloramphenicol inhibited growth to the same extent as kanamycin at high concentrations. Ampicillin did not completely inhibit growth. B. Spectinomycin, chloramphenicol, and kanamycin were found to inhibit growth.

### SUPPLEMENTARY TABLES

#### Supplementary Table 1: openCIDAR Broad Host Range MoClo Destination Vectors and Fluorescent Reporters

| Plasmid name | CIDAR MoClo Level | Resistance | Addgene ID |
|---|---|---|---|
| pBBR1_DVK_AE | 1 | Kanamycin | 188946 |
| pBBR1_DVK_EF | 1 | Kanamycin | 188947 |
| pBBR1_DVK_FG | 1 | Kanamycin | 188948 |
| pBBR1_DVK_GH | 1 | Kanamycin | 188949 |
| pBBR1_DVS_AF | 2 | Spectinomycin | 188950 |
| pBBR1_DVS_AG | 2 | Spectinomycin | 188951 |
| pBBR1_DVS_AH | 2 | Spectinomycin | 188952 |
| pBBR1_DVC_AF | 2 | Chloramphenicol | 188953 |
| pBBR1_DVC_AG | 2 | Chloramphenicol | 188954 |
| pBBR1_DVC_AH | 2 | Chloramphenicol | 188955 |
| pBBR1_DVK_AE(J23100-B0032m-E0040m-B0015) | 1 | Kanamycin | 197657 |
| pBBR1_DVK_AE(J23100-B0033m-E0040m-B0015) | 1 | Kanamycin | 197658 |
| pBBR1_DVK_AE(J23100-B0034m-E0040m-B0015) | 1 | Kanamycin | 197659 |
| pBBR1_DVK_AE(J23100-BCD2-E0040m-B0015) | 1 | Kanamycin | 197660 |
| pBBR1_DVK_AE(J23100-RiboJ-BCD2-E0040m-B0015) | 1 | Kanamycin | 197661 |
| pBBR1_DVK_AE(J23100-RiboJ-UTR1-E0040m-B0015) | 1 | Kanamycin | 197662 |
| pBBR1_DVK_AE(J23103-B0032m-E0040m-B0015) | 1 | Kanamycin | 197663 |
| pBBR1_DVK_AE(J23103-B0033m-E0040m-B0015) | 1 | Kanamycin | 197664 |
| pBBR1_DVK_AE(J23103-B0034m-E0040m-B0015) | 1 | Kanamycin | 197665 |
| pBBR1_DVK_AE(J23103-BCD2-E0040m-B0015) | 1 | Kanamycin | 197666 |
| pBBR1_DVK_AE(J23103-RiboJ-BCD2-E0040m-B0015) | 1 | Kanamycin | 197667 |
| pBBR1_DVK_AE(J23103-RiboJ-UTR1-E0040m-B0015) | 1 | Kanamycin | 197668 |
| pBBR1_DVK_AE(J23106-B0032m-E0040m-B0015) | 1 | Kanamycin | 197669 |
| pBBR1_DVK_AE(J23106-B0033m-E0040m-B0015) | 1 | Kanamycin | 197670 |
| pBBR1_DVK_AE(J23106-B0034m-E0040m-B0015) | 1 | Kanamycin | 197671 |
| pBBR1_DVK_AE(J23106-BCD2-E0040m-B0015) | 1 | Kanamycin | 197672 |
| pBBR1_DVK_AE(J23106-RiboJ-BCD2-E0040m-B0015) | 1 | Kanamycin | 197673 |
| pBBR1_DVK_AE(J23106-RiboJ-UTR1-E0040m-B0015) | 1 | Kanamycin | 197674 |

**Supplementary Table 2: Known hosts for pBBR1 and its derivatives**

| Organism | NCBI TaxID | Class | Reference |
|---|---|---|---|
| Komagataeibacter nataicola | NCBI:txid265960 | Alphaproteobacteria | This study |
| Komagataeibacter rhaeticus | NCBI:txid215221 | Alphaproteobacteria | Florea et al. 2016 |
| Komagataeibacter xylinus | NCBI:txid28448 | Alphaproteobacteria | Kovach et al. 1995 |
| Novacetimonas hansenii | NCBI:txid436 | Alphaproteobacteria | Florea et al. 2016 |
| Cupriavidus necator | NCBI:txid106590 | Betaproteobacteria | Kovach et al. 1995 |
| Bartonella bacilliformis | NCBI:txid774 | Alphaproteobacteria | Kovach et al. 1995 |
| Bordetella pertussis | NCBI:txid520 | Betaproteobacteria | Antoine and Locht 1992 |
| Bordetella bronchiseptica | NCBI:txid518 | Betaproteobacteria | Antoine and Locht 1992 |
| Brucella spp. | NCBI:txid234 | Alphaproteobacteria | Kovach et al. 1994 |
| Caulobacter vibrioides | NCBI:txid155892 | Alphaproteobacteria | Kovach et al. 1995 |
| Escherichia coli | NCBI:txid562 | Gammaproteobacteria | Antoine and Locht 1992 |
| Paracoccus denitrificans | NCBI:txid266 | Alphaproteobacteria | Kovach et al. 1995 |
| Pseudomonas fluorescens | NCBI:txid294 | Gammaproteobacteria | Kovach et al. 1995 |
| Pseudomonas putida | NCBI:txid303 | Gammaproteobacteria | Antoine and Locht 1992 |
| Sinorhizobium meliloti | NCBI:txid382 | Alphaproteobacteria | Antoine and Locht 1992 |
| Rhizobium leguminosarum bv. Viciae | NCBI:txid387 | Alphaproteobacteria | Kovach et al. 1995 |
| Cereibacter sphaeroides | NCBI:txid1063 | Alphaproteobacteria | Kovach et al. 1995 |
| Salmonella enterica subsp. Enterica serovar Typhimurium | NCBI:txid90371 | Gammaproteobacteria | Kovach et al. 1995 |
| Vibrio cholerae | NCBI:txid666 | Gammaproteobacteria | Antoine and Locht 1992 |
| Xanthomonas campestris | NCBI:txid339 | Gammaproteobacteria | Kovach et al. 1995 |
| Paraburkholderia sacchari | NCBI:txid159450 | Betaproteobacteria | Guamán et al. 18 |
| Chromobacterium violaceum | NCBI:txid536 | Betaproteobacteria | Liow et al. 20 |
| Vibrio natriegens | NCBI:txid691 | Gammaproteobacteria | Tschirhart et al. 19 |
| Rhodopseudomonas palustris | NCBI:txid1076 | Alphaproteobacteria | Immethun et al. 22 |
| Shewanella oneidensis | NCBI:txid70863 | Gammaproteobacteria | Suzuki et al. 20 |
| Burkholderia cepacia | NCBI:txid292 | Betaproteobacteria | Lefebre and Valvano 02 |
| Gluconobacter oxydans | NCBI:txid442 | Alphaproteobacteria | Fricke et al. 20 |
| Agrobacterium tumefaciens | NCBI:txid358 | Alphaproteobacteria | Yamamoto et al. 18 |
| Agrobacterium rhizogenes | NCBI:txid359 | Alphaproteobacteria | Yamamoto et al. 18 |
| Agrobacterium vitis | NCBI:txid373 | Alphaproteobacteria | Yamamoto et al. 18 |
| Geobacter sulfurreducens | NCBI:txid35554 | Deltaproteobacteria | Chan et al. 15 |
| Methylococcus capsulatus | NCBI:txid414 | Gammaproteobacteria | Tapscott el al. 19 |
| Cupriavidus metallidurans | NCBI:txid119219 | Betaproteobacteria | Trepreau et al. 14 |

**Supplementary Table 3: Primers and probes for plasmid copy number quantification**

| Organism | Target | Primer name | Type | Conjugation | Sequence |
|---|---|---|---|---|---|
| *E.coli* | cysG | 1347 | F | N/A | AGCCATTACTGAAACGACC |
| *E.coli* | cysG | 1348 | R | N/A | GCTGAATTTGTTGCAGTCC |
| *E.coli* | cysG | 1349 | P | FAM / ZEN Iowa Black FQ | ACCAACCAGCACCACTTCACCG |
| *P.putida* | ileS | 1350 | F | N/A | GGACAACCCATACAAGACC |
| *P.putida* | ileS | 1351 | R | N/A | TCAAAGCACCAGTTCACC |
| *P.putida* | ileS | 1352 | P | FAM / ZEN Iowa Black FQ | TCCGCGCCCTGGCCGA |
| *C.necator* | panC | 1775 | F | N/A | CATTTCCTCCATCCAAGAGC |
| *C.necator* | panC | 1776 | R | N/A | GGGTACTTGTCGAAATCCTC |
| *C.necator* | panC | 1777 | P | FAM / ZEN Iowa Black FQ | ACCGCGCTGCCTTCGTGCCC |
| *K.nataicola* | cysG | 1778 | F | N/A | CTCGGGTATGGTGCTTTC |
| *K.nataicola* | cysG | 1779 | R | N/A | GCATGAACATTCCCGTCT |
| *K.nataicola* | cysG | 1780 | P | FAM / ZEN Iowa Black FQ | TGCGCACCAGCCCGCGATGG |
| All | BBR1 *rep** | 1353 | F | N/A | TGGTCAGCCAGAAAACACTT |
| All | BBR1 *rep** | 1354 | R | N/A | GTCCTTGACTGCGTATTGGA |
| All | BBR1 *rep** | 1355 | P | HEX / ZEN Iowa Black FQ | AGCTCATCGGACGTTCTTTGCGGACGG |
